# BET inhibition disrupts transcription but retains enhancer-promoter contact

**DOI:** 10.1101/848325

**Authors:** Nicholas T. Crump, Erica Ballabio, Laura Godfrey, Ross Thorne, Emmanouela Repapi, Jon Kerry, Marta Tapia, Peng Hua, Panagis Filippakopoulos, James O. J. Davies, Thomas A. Milne

## Abstract

In higher eukaryotes, enhancers are DNA sequences that enable complex temporal and tissue-specific regulation of genes. Although it is not entirely clear how enhancer-promoter interactions can increase gene expression, this proximity has been observed in multiple systems at multiple loci and is thought to be essential for the maintenance of gene expression. The formation of phase condensates is thought to be an essential component of enhancer function. Here, we show that pharmacological targeting of cells with inhibitors of BET (Bromodomain and Extra-Terminal domain) proteins can have a strong impact on transcription but very little impact on enhancer-promoter interactions. Treatment with 1,6-hexanediol, which dissolves phase condensate structures and reduces BET and Mediator protein binding at enhancers, can also have a strong effect on gene transcription, without disrupting enhancer-promoter interactions. These results suggest that activation of transcription and maintenance of enhancer-promoter interactions are separable events. Our findings further suggest that enhancer-promoter interactions are not dependent on high levels of BRD4 (Bromodomain-containing protein 4) and Mediator, and are likely maintained by a complex set of factors including additional activator complexes and loop extrusion by CTCF/cohesin.

## Introduction

In higher eukaryotes, enhancers are DNA sequences that allow for the complex regulation of genes in different tissues and at different times ^1^. Despite the importance of enhancers, very little is known about exactly how they function, although they have been proposed to act mainly as binding platforms for the assembly of protein complexes that can promote gene activation ^2,3^. A key aspect to this assembly is the binding of sequence-specific DNA binding factors such as transcription factors (TFs). Enhancers can be situated far away from the genes they regulate ^1,4^. Although not always the case ^5,6^, at many gene loci proximity between enhancers and promoters is thought to be essential for enhancer function and gene activation ^7,8^. How these enhancer-promoter interactions are initiated and maintained is not clearly understood.

Emerging work suggests that enhancers function within larger domains, the boundaries of which are defined by the combined effects of CTCF-marked boundary regions and cohesin looping, through a process known as loop extrusion ^9,10^. It is not entirely clear how these higher order structures impact enhancer function, but generally speaking functional enhancer-promoter interactions are limited to genes within or at the edges of domains. The genome-wide existence of these more localized enhancer-promoter looping structures has been demonstrated by global chromosome conformation capture (3C) techniques ^11,12^. However, unless each sample is sequenced extremely deeply ^13^ (something that is not practical for most experiments), Hi-C is not able to delineate enhancer-promoter interactions at high resolution, so it is difficult to study these structures in detail genome-wide. The high complexity associated with Hi-C libraries has meant that there have been multiple attempts to develop high-throughput methods to provide more information specifically about enhancer-promoter interactions ^11,12,14,15^, but the highest resolution studies focus on individual genes/enhancers using next generation techniques such as 4C ^16^, UMI-4C seq ^17^, Next Generation Capture-C ^18^, Tri-C ^19^ and tiled Capture-C ^20^.

Whilst loop extrusion mediated by CTCF and cohesin is the most common explanation for controlling large scale chromatin structure, less is known about what stabilizes more localized enhancer-promoter loops. Possible models include CTCF/cohesin-stabilized enhancer-promoter interactions ^21–23^ and protein/RNA complexes bound to both the enhancer and promoter that interact with one another ^24–26^. Binding of these complexes is likely initiated by key sequence-specific TFs. The presence of specific histone modifications at the enhancer is thought to contribute to one or all of these models, mainly by stabilizing the presence of specific protein complexes, such as cohesin binding to H3K4me1 or BRD4 (Bromodomain-containing protein 4) binding to H3K27ac ^27–29^. Recent work from our lab also suggests that at H3K79me2/3-dependent enhancer elements (KEEs), the presence of H3K79me2/3 can help maintain open chromatin regions to facilitate the binding of sequence specific transcription factors, and is required for enhancer-promoter interaction ^30^. This could constitute a more general principal where histone modifications help regulate DNA accessibility and TF binding, and ultimately the formation of enhancer-promoter loops.

Super-enhancers (SEs) are enhancers with enriched levels of binding of TFs and activating complexes such as BRD4 and Mediator and are associated with high levels of transcriptional activity ^2,31^. In cancer cells, important oncogenes are often associated with super-enhancers ^32,33^. Recent work has shown that many enhancer-associated factors, such as Mediator (e.g. MED1) and BRD4, assemble into phase-separated activation complexes, and these interactions appear to be integral to their ability to activate transcription ^2,3,34–37^. Further, several studies have linked the phase separation-mediated assembly of activation complexes at enhancers (particularly SEs) and promoters to the initiation and maintenance of interactions between enhancers and promoters ^2,3,35–37^.

Taken together, these various strands of evidence suggest the following model: a) loop extrusion via CTCF/cohesin complexes generates higher-order chromatin structures; b) TFs bound to enhancers and promoters assemble phase condensates made up of chromatin proteins such as BRD4 and coactivators such as MED1; c) histone modifications maintain accessibility for the binding of TFs and create additional affinities to further stabilize complexes; d) the CTCF/cohesin-delimited structures create a smaller DNA compartment, increasing the frequency of random interactions between complexes bound at enhancers and promoters; and e) the phase-separated condensates containing BRD4 and Mediator anchored at the enhancer and promoter act as a bridge to stabilize these enhancer-promoter interactions. Thus, the model posits that formation of phase condensates is a key requirement for at least a subset enhancer-promoter looping. Recent work testing this model indicated no loss of enhancer-promoter contact following degron-mediated loss of MED14, suggesting that Mediator is not responsible for these interactions ^21^. However, this study used Hi-C (binned at 5 kb) and promoter capture Hi-C, which are relatively low resolution and low sensitivity techniques, so it is possible that perturbations in specific enhancer-promoter contacts may have been missed. It therefore remains unclear whether BRD4 and Mediator play any role in organizing chromatin structure and maintaining enhancer-promoter interactions at active genes.

In this paper, we directly test the role of BRD4 and Mediator in enhancer-promoter interactions by performing high resolution Next Generation Capture-C ^18^ at over 60 specific loci in cells treated with BET inhibitors and 1,6-hexanediol. This technique provides the greatest resolution and sensitivity of all the available 3C methods for higher eukaryotic cells ^38^. Data are generated using four-cutter restriction enzymes and are of sufficient sequencing depth that they can be reported at single restriction fragment resolution. In addition, the method is highly sensitive and reproducible, meaning that changes in interaction frequency can be analyzed quantitatively under different conditions at many genes simultaneously.

We find that reduction of BRD4 and Mediator binding at enhancers has a dramatic and rapid effect on gene expression, but enhancer-promoter looping structures remain stably intact. This suggests that the function of these activation complexes at enhancers does not involve stabilization of the enhancer-promoter interaction. Instead, we see evidence of CTCF and cohesin binding at many enhancers, indicating that these complexes can stabilize and maintain looping structures even in the presence of reduced transcription and activation complexes at the enhancer. Finally, our results demonstrate that stabilization of enhancer-promoter interactions and promotion of transcription are separable events, and that the presence of an enhancer-promoter loop is not sufficient for the maintenance of transcription.

## Results

### BET and Mediator proteins bind to active enhancers and promoters in leukemia cells

BRD4 and Mediator binding are key characteristics of enhancers, particularly super-enhancers, which are defined as having high levels of these proteins over extended regions ^31–33^. We analyzed levels of BET-domain (Bromodomain and Extra-Terminal domain) proteins and Mediator subunits at ATAC peaks genome-wide in the leukemia cell line SEM, with peaks ranked by the relative levels of H3K4me3 and H3K4me1, thereby separating promoter and enhancer loci (Fig 1a). BET domain proteins (i.e. BRD2, BRD3 and BRD4) and Mediator subunits MED1, MED12 and MED26 all showed an enrichment at both promoter and enhancer ATAC peaks, comparable to the distribution of H3K27ac (Fig 1a, Supplementary Fig 1a). Consistent with the idea that BRD4 physically interacts with Mediator ^39–42^, BRD4 binding positively correlated with all three Mediator subunits at ATAC peaks (Fig 1b). In contrast, although they appear to overlap at a subset of loci (Fig 1a), CTCF and the cohesin subunit RAD21 clustered separately from BRD4/Mediator (Fig 1b), showing a distinct distribution at ATAC peaks, with similar levels at promoters, enhancers and other accessible regions (Fig 1a, Supplementary Fig 1a).

**Figure 1.**
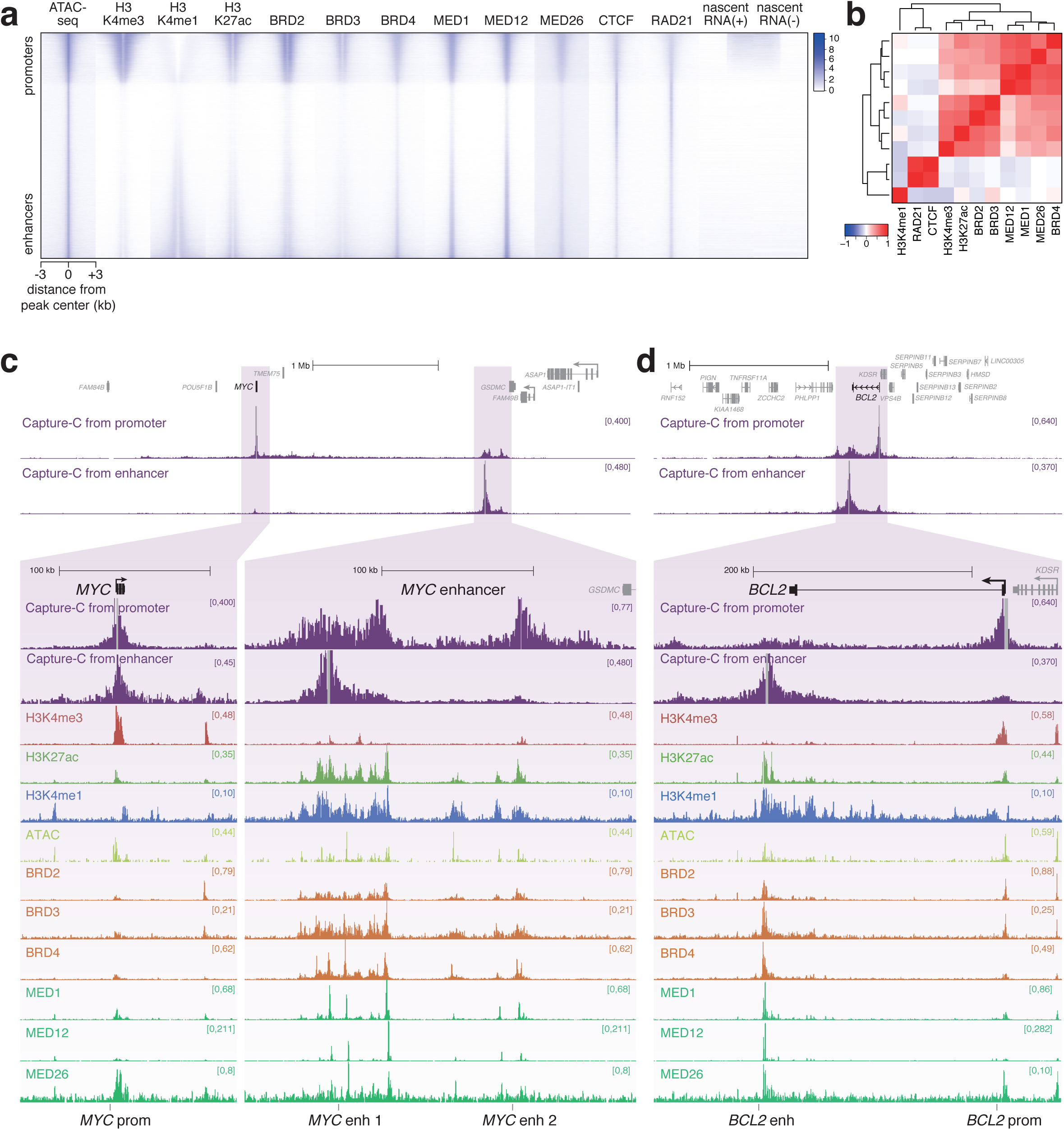
BET proteins and Mediator are a key feature of enhancers. **a** Heatmap comparing levels of histone modifications, chromatin proteins and stranded nascent RNA-seq at ATAC-seq peaks in SEM cells. Peaks are ranked based on the relative levels of H3K4me3 and H3K4me1, placing promoter-like ATAC-seq peaks towards the top and enhancer-like ATAC peaks towards the bottom. **b** Pearson correlation coefficients for ChIP-seq data at ATAC-seq peaks shown in (a). Dendrogram shows hierarchical clustering of datasets. **c** Capture-C, ChIP-seq and ATAC-seq at the *MYC* gene and enhancer region in SEM cells. Capture-C was conducted using the *MYC* promoter or enhancer region as the viewpoint, indicated by vertical gray bars, and is displayed as the mean of three biological replicates. Locations of primers used for BRD4/Mediator ChIP-qPCR in Fig 4 and Supplementary Fig 2 are shown at the bottom of the figure. **d** Capture-C, ChIP-seq and ATAC-seq data at *BCL2*, as in (c).

Since BRD4 is associated with the enhancers and promoters of highly transcribed genes (Fig 1a and Supplementary Fig 1b, c) ^31–33^, we wanted to examine its role in enhancer function in more detail. Two classic BRD4-dependent genes are *MYC* and *BCL2* ^43,44^. We used the high resolution 3C technology Next Generation Capture-C ^18^ to identify enhancers for these genes based on their ability to interact with their promoters (Fig 1c, d) ^30^. Our Capture-C experiments in SEM cells revealed a high frequency of interaction between the *MYC* promoter and a large (~200 kb) region, composed of two major domains, located ~1.7 Mb away. This long-distance interaction has also been observed and characterized in several other cell types ^22,23,45,46^. Reciprocal Capture-C from the more proximal of the two enhancer regions demonstrated contact exclusively with the *MYC* promoter, avoiding intervening regions, as well as a relatively weak interaction with the more distal enhancer domain (Fig 1c). The enhancer is marked with broad domains of H3K27ac, BET proteins and Mediator, with much higher levels than at the *MYC* promoter (Fig 1c, lower). In addition, we observed multiple peaks of chromatin accessibility by ATAC-seq (Fig 1c), and enrichment for multiple transcription factors (Supplementary Fig 1e). These characteristics are consistent with the region acting as a strong enhancer to regulate *MYC* expression; indeed, it is defined as a super-enhancer following established criteria ^31–33^.

We and others have previously demonstrated the presence of an enhancer at the 3’ end of *BCL2* in SEM cells ^30,47,48^, and Capture-C from the enhancer illustrates its interaction with the *BCL2* promoter (Fig 1d). As at *MYC*, this region is identified as a super-enhancer and is marked by elevated levels of H3K27ac, BRD4 and Mediator, relative to the *BCL2* promoter (Fig 1d, lower).

To investigate the association of BRD4 with enhancer-promoter interactions on a larger scale, we analyzed Capture-C data for the promoters of 62 genes (Supplementary Table 1). We used ChIP-seq data for a number of enhancer-associated features to ask whether these features were commonly associated with an increased frequency of interaction with gene promoters. Indeed, H3K4me1, H3K27ac, BRD4 and MED1 were all associated with a higher frequency of interaction with promoters, compared to the average interaction frequency across the interaction domains (Supplementary Fig 1d). In contrast, the repressive histone modification H3K27me3 was found at loci with reduced promoter contact frequency (Supplementary Fig 1d). This analysis revealed that BRD4 and Mediator binding is associated with a higher frequency of interaction with promoters, potentially implicating these proteins in stabilizing enhancer-promoter contacts at these and potentially other genes genome-wide.

### Loss of BET and Mediator binding is associated with large transcriptional changes at key oncogenic gene targets

In order to investigate the role of BRD4 in enhancer function, we used the small molecule inhibitor IBET-151, which disrupts binding of BET protein bromodomains to acetyllysine residues. IBET is known to disrupt transcription, so we wanted to use a short treatment time to limit secondary effects from regulatory events downstream of initial transcriptional changes. We used qRT-PCR to assess how quickly IBET treatment affects gene expression. However, the stability of mature transcripts means that there is often a delay between decreased transcription and changes in mRNA levels (Supplementary Fig 2a). We therefore used primers against intronic sequences to quantify levels of the more labile pre-mRNA. Strikingly, we observed very rapid changes in transcription, with levels of *MYC* pre-mRNA decreasing after only 15 min IBET treatment (Fig 2a, *left*). Levels of *BCL2* were also sensitive to IBET treatment, with ~50 % loss after 90 min (Fig 2a). In contrast, a comparable decrease in mature *BCL2* mRNA was not detected before 3h (Supplementary Fig 2a). IBET also resulted in the similarly rapid upregulation of a number of genes (Fig 2a, *right*).

**Figure 2.**
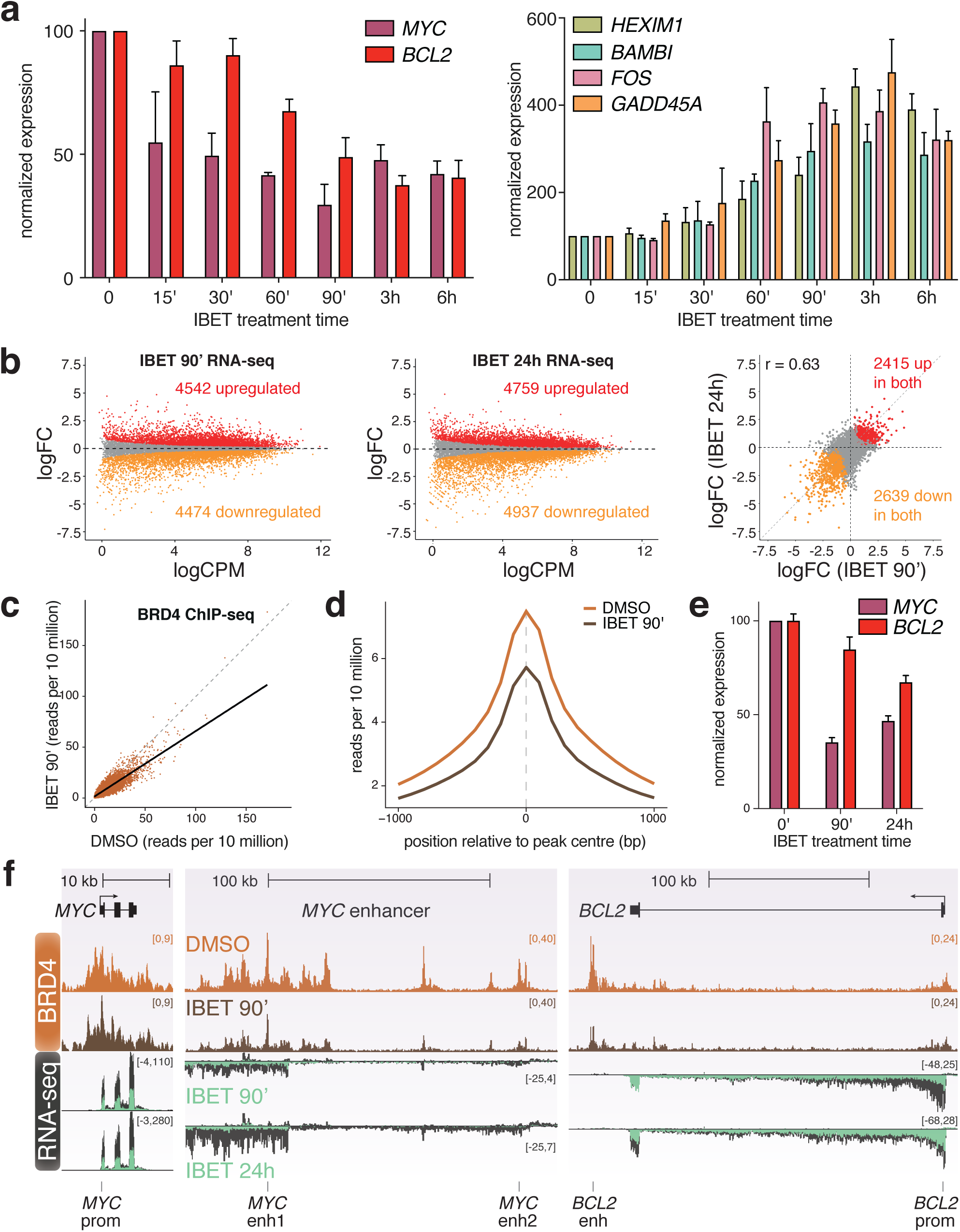
IBET treatment results in large-scale transcriptional changes. **a** *Left*: qRT-PCR analysis of gene expression following IBET treatment for the indicated times, using intronic PCR primers. *Right*: qRT-PCR analysis of gene expression using mature mRNA PCR primers. Values are normalized to *YWHAZ* mature mRNA levels, relative to DMSO treatment. Mean of three biological replicates; error bars show SEM. **b** MA plots for changes in nascent RNA levels following 90 min (*left*) or 24h (*middle*) treatment with 1 µM IBET. *Right*: correlation of log fold-change (logFC) of gene expression following IBET treatment for 90 min or 24h. Statistically significant differences (red: increased; orange: decreased; gray: unchanged) from three biological replicates, FDR <0.05. **c** Correlation of BRD4 ChIP-seq reads at BRD4 peaks from SEM cells treated with DMSO or 1 µM IBET for 90 min. **d** Metaplot of mean BRD4 levels at BRD4 peaks in SEM cells treated with DMSO or 1 µM IBET for 90 min. **e** Quantification of *MYC* and *BCL2* nascent RNA-seq levels in SEM cells treated with DMSO or 1 µM IBET for 90 min or 24h. Mean of three biological replicates, normalized to expression in DMSO; error bars show SEM. **f** BRD4 ChIP-seq and nascent RNA-seq at the *MYC* gene and enhancer and at *BCL2*. BRD4 ChIP-seq was conducted on SEM cells treated with DMSO or 1 µM IBET for 90 min. Stranded nascent RNA-seq tracks are an overlay of data from cells treated with DMSO (gray) or IBET (green) for 90 min (above) or 24 h (below), representative of three biological replicates.

To assess the direct effects of BRD4 inhibition, we chose a 90 min IBET treatment time, based on the qRT-PCR data. We analyzed the global transcriptional response to IBET by sequencing nascent RNA ^49^, which provides a much more direct measure of transcriptional output compared to steady state RNA-seq (Fig 2b, Supplementary Table 2). As a comparison, we also sequenced nascent RNA following 24h IBET treatment (Fig 2b, Supplementary Table 2). The number of differentially-expressed genes was comparable after 90 min and 24h IBET treatment, and there was reasonable correlation between the intensity of the changes under each condition (R = 0.63, Fig 2b, *right*), suggesting that the shorter treatment time is sufficient to capture the immediate effects of BET domain inhibition.

Surprisingly, given the role of BRD4 in promoting transcription, we observed similar numbers of up- and downregulated genes, several of which we confirmed by qRT-PCR (Fig 2a, Supplementary Fig 2b). This may be explained by the observation that BRD4 has a role in gene repression as well as activation ^50,51^, suggesting that some of the upregulated genes may be direct targets of BRD4. In addition, upregulated genes were enriched for biological pathways associated with a response to chemical stimulus (Supplementary Fig 2c) indicating that there may also be an indirect response to drug treatment, consistent with previously published work ^52^.

We confirmed the genome-wide reduction of BRD4 binding after 90 min incubation with IBET by ChIP-seq (Fig 2c, d) and ChIP-qPCR (Supplementary Fig 2d). At the *MYC* and *BCL2* enhancers, BET inhibition was associated with reduced transcription as well as a decrease in BRD4 binding (Fig 2e, f, Supplementary Fig 2d). Consistent with the loss of MED1 foci in cells following BET inhibition ^34^, treatment with IBET also resulted in the dissociation of Mediator subunits from chromatin, with reductions in MED1 and MED12 binding at the *MYC* enhancer (Supplementary Fig 2d). Thus, IBET treatment reduces the level of BRD4 and Mediator binding to enhancers, and this is associated with a reduction in transcription.

### Loss of BET and Mediator binding has very little effect on enhancer-promoter looping

BRD4 and MED1 have recently been shown to be present in phase condensate clusters in the nucleus ^2,34,35^, and these structures are proposed to be important for the function of super-enhancers, potentially by mediating interactions with target gene promoters. In previously published work, treatment of cells with the small molecule inhibitor JQ1, which, like IBET-151, disrupts binding of the BRD4 BET domain to acetyllysine residues, prevented clustering of BRD4 and MED1 ^34^, indicating that association with chromatin is integral to phase condensation. We therefore asked whether inhibition of BRD4 binding would disrupt enhancer-promoter interactions, as this might explain the strong effect of IBET treatment on transcription.

Strikingly, however, treatment with IBET had little or no effect on enhancer-promoter association. At the *MYC* enhancer, the major regions of contact with the promoter remained virtually unchanged by 90 min IBET treatment, with only small differences in interaction frequency (Fig 3a). A similar result was observed in the reciprocal analysis from the enhancer, demonstrating only a subtle increase in interactions at the promoter (Fig 3a, *below*). Even at a later 24h timepoint, these enhancer-promoter interactions were mostly retained (Fig 3a), suggesting that the looping structure is stable in the absence of high levels of BRD4. Whilst we do observe some rearrangement of the enhancer-promoter interactions after 24h IBET treatment, with an apparent shift from the distal to the more proximal region of the enhancer (Fig 3a), the broad interaction profile is maintained. It is worth noting that these changes do not correlate with the early disruption of gene expression, as transcription of *MYC* was decreased after only 15 min IBET treatment and remained inhibited at 24h (Fig 2a, e). We observed a similar maintenance of enhancer-promoter interactions at *BCL2*, where contact between the promoter and enhancer was preserved after even 24h IBET treatment (Fig 3b) despite a decrease in transcription (Fig 2a, e), arguing that although reduced BRD4 and Mediator can impact gene expression, maintaining high levels of these factors is not required for enhancer-promoter interactions at *MYC* or *BCL2*.

**Figure 3.**
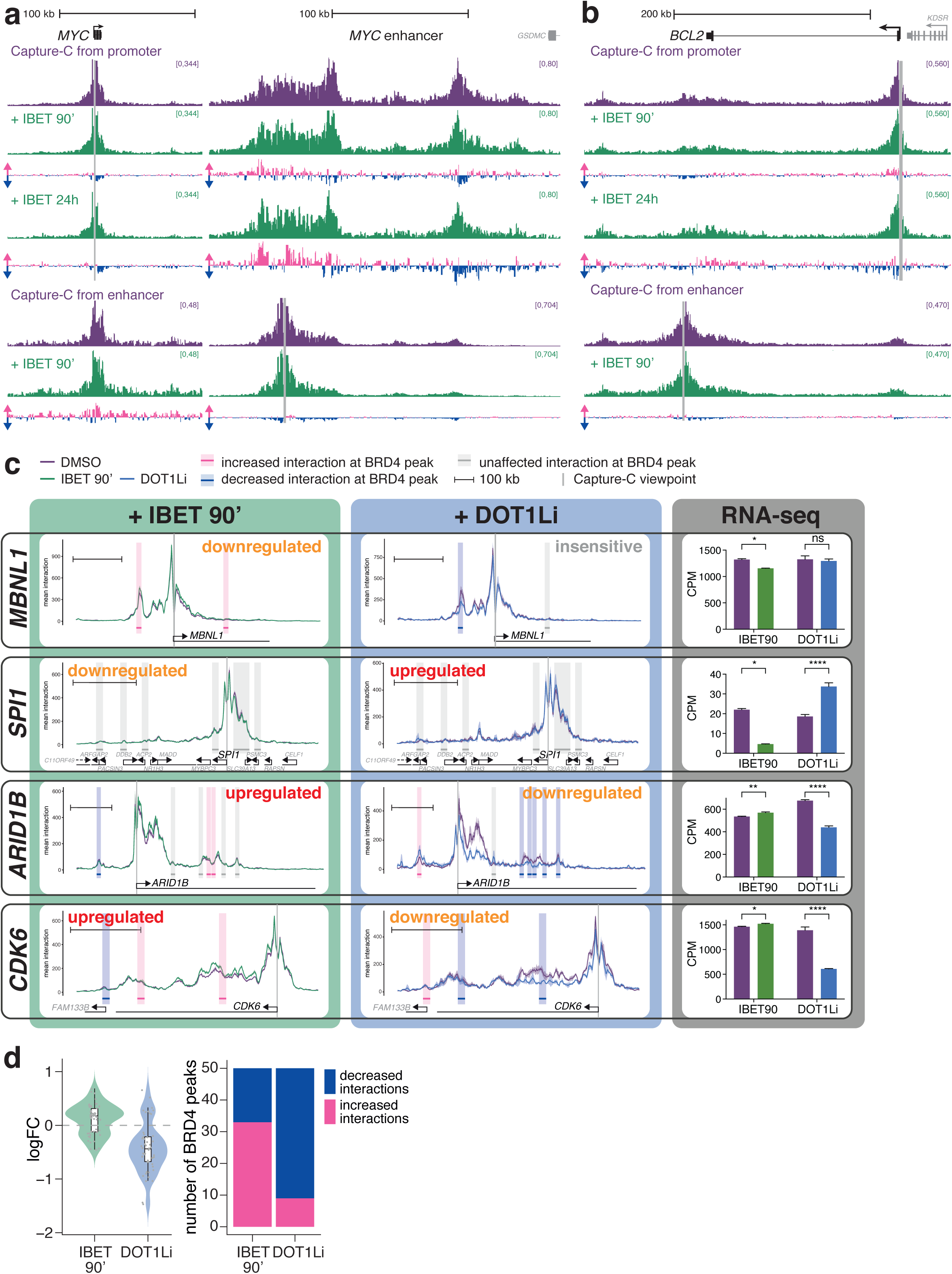
IBET treatment has minimal effects on enhancer-promoter interactions. **a** Capture-C from the *MYC* promoter (*above*) or enhancer (*below*) following 90 min DMSO treatment (purple) or 90 min or 24h 1 µM IBET treatment (green). Only the promoter and enhancer regions are shown. Differential tracks show the change in profile in IBET-treated samples compared to DMSO treatment for the same time period: pink bars show increases; blue bars show decreases. Mean of three biological replicates. **b** Capture-C from the *BCL2* promoter (*above*) or enhancer (*below*), as in (a). **c** Capture-C traces at genes following treatment with DMSO (purple line), IBET for 90 min (*left*, green line) or EPZ-5676 for 7d (DOT1Li, *middle*, blue line). Ribbon shows +/−1 SD for three replicates. Vertical gray bar indicates the capture point for each gene. Horizontal bars show 10 kb region around BRD4 ChIP-seq peaks. Shading highlights effect of IBET treatment on promoter interaction frequency within that window: pink bars indicate statistically-significant increases; blue bars indicate decreases; gray bars indicate no significant difference (Holm-Bonferroni adjusted p-value <0.05, paired Mann-Whitney test). Scale bars show 100 kb. Transcriptional effect of the drug treatment on the gene is indicated. *Right*: Nascent RNA-seq levels for each gene under control or drug treatment conditions. **** = FDR <0.0001, * = FDR <0.05, ns = no significant change. DOT1Li Capture-C and RNA-seq data are taken from^30^. **d** *Left*: change in interaction frequency (mean logFC of three replicates) between promoters and BRD4 peaks (10 kb windows) following 90 min IBET or 7 day DOT1Li treatment. Boxplot shows median and IQR. Dots represent individual BRD4 peaks. *Right*: number of BRD4 peaks (10 kb windows) that show statistically-significant (Holm-Bonferroni adjusted p-value <0.05, paired Mann-Whitney test) increases (pink) or decreases (blue) following 90 min IBET or 7 day DOT1Li treatment.

A more widespread analysis of Capture-C at a further 60 genes (Supplementary Table 1) demonstrated a similar response to IBET treatment, with minimal changes in promoter contacts (Fig 3c, Supplementary Fig 3a), whether they were up- or downregulated or transcriptionally unaffected by treatment (Supplementary Fig 3b). This is in striking contrast to previous work from our lab showing a strong correlation between loss of enhancer-promoter interactions and reduction of transcription following DOT1L inhibition (DOT1Li), which results in decreased activity at H3K79me2/3-marked enhancers ^30^ (see Fig 3c for comparison). Surprisingly, IBET treatment led to small increases in interaction frequency at a number of genes (Fig 3c, Supplementary Fig 3a), despite the loss of BRD4. In some cases, this correlated with a slight upregulation of transcription (e.g. *CDK6*, Fig 3c), but in other cases it correlated with downregulation (e.g. *MBNL1*, Fig 3c). In order to quantify these differences, we measured the changes in interaction frequency at BRD4 peaks, reasoning that these sites were most likely to be affected by loss of BRD4 binding. Because of the broad nature of the interaction profile, we used a 10 kb window centered on each peak (highlighted regions in Fig 3c, Supplementary Fig 3a). The majority of loci that showed statistical changes revealed a slight increase in interaction frequency following IBET treatment (Fig 3d; mean logFC=0.11). This lack of a strong effect is not due to a limitation of the Capture-C technique, as DOT1Li-treated cells demonstrated a clear reduction in interaction frequency (Fig 3c, d, mean logFC=−0.42).

Longer treatment with IBET did not result in the delayed disruption of enhancer-promoter interactions (Supplementary Fig 3c). Indeed, there was a clear correlation between the Capture-C changes observed at BRD4 peaks following 90 min or 24h treatment with IBET (Supplementary Fig 3d, R=0.62), demonstrating that these chromatin structures are stable in the absence of BRD4 and Mediator. Treatment with another BET domain inhibitor, JQ1, which also disrupts the chromatin association of BRD4 (Supplementary Fig 4a), resulted in similarly subtle effects on promoter interaction profiles (Supplementary Fig 4b). Taken together, these results argue that whilst high levels of BRD4/Mediator binding may be required for enhancer function and gene transcription, they do not act primarily by stabilizing physical contact with the gene promoter.

Whilst IBET treatment results in the dissociation of BRD4 from chromatin, it is possible that other factors remain at enhancers that could facilitate promoter contact via low affinity clustering interactions. It is also possible that the lower levels of BRD4 and MED1 binding that remain at enhancers after IBET treatment are sufficient to maintain enhancer-promoter interactions, despite the disruption of transcription. To induce a stronger, more generalized effect at these loci, we used 1,6-hexanediol, which is commonly employed to dissolve phase condensates, structures that have recently been reported to be important for super-enhancer function ^2,53,54^. Hexanediol treatment had a striking effect on BRD4 binding at the *MYC* and *BCL2* enhancers (Fig 4a), and MED1 levels were also reduced at the *MYC* enhancer (Fig 4a). Hexanediol resulted in rapid changes in gene expression by nascent RNA-seq, with differential expression of more than 8000 genes after only 30 min (Fig 4b, Supplementary Table 2).

**Figure 4.**
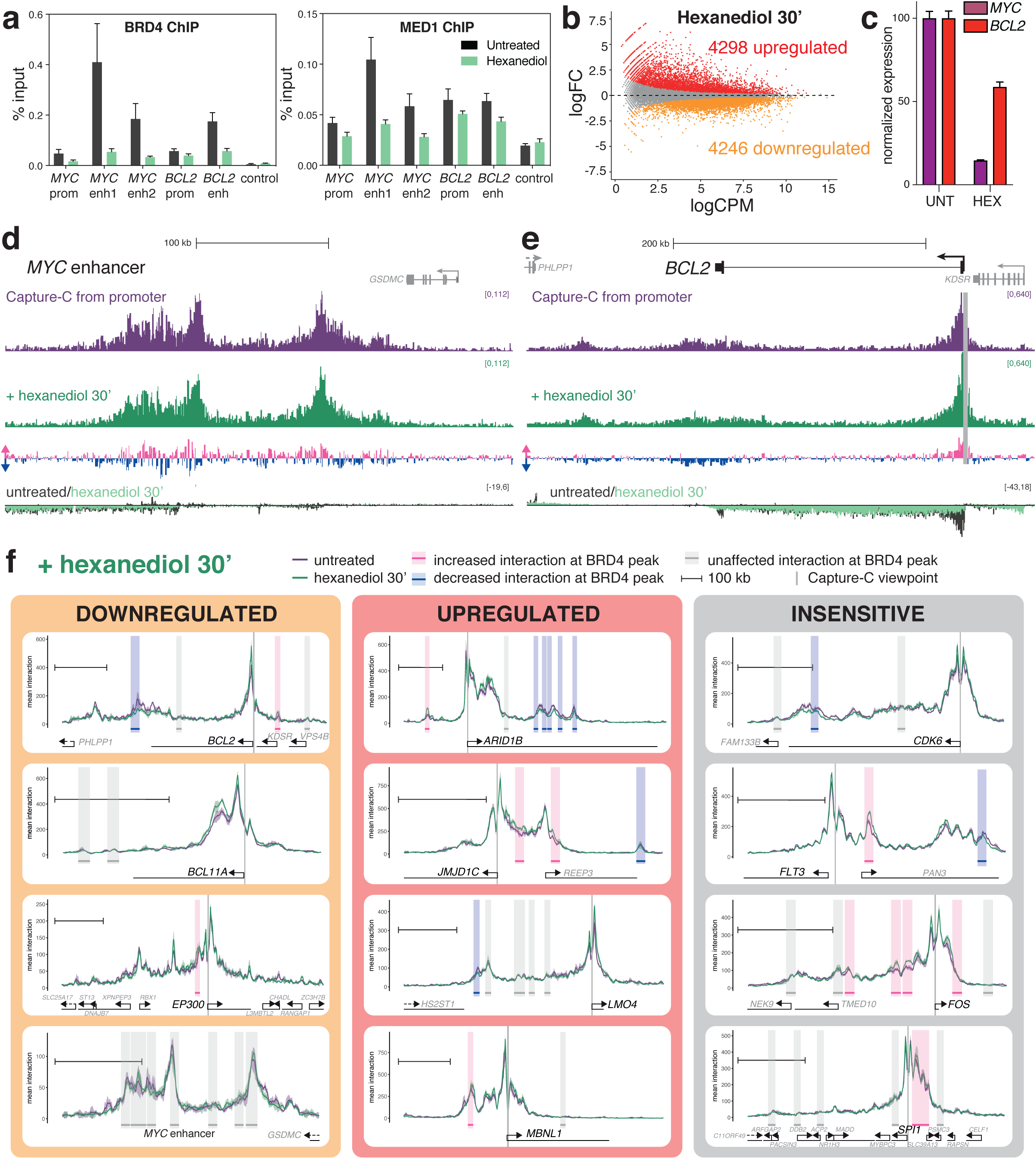
Dissolution of phase condensate structures with 1,6-hexanediol does not perturb enhancer-promoter interactions. **a** ChIP-qPCR for BRD4 and MED1 from untreated SEM cells (black) or following 30 min treatment with 1.5 % 1,6-hexanediol (green). Mean of four biological replicates; error bars show SEM. **b** MA plot for changes in nascent RNA levels following 30 min treatment with 1.5 % 1,6-hexanediol. Mean of three biological replicates. Statistically significant differences (red: increased; orange: decreased; gray: unchanged) from three biological replicates, FDR <0.05. **c** Quantification of *MYC* and *BCL2* nascent RNA-seq levels in untreated SEM cells or cells treated with 1.5 % 1,6-hexanediol for 30 min. Mean of three biological replicates, normalized to expression in untreated cells; error bars show SEM. **d** Capture-C from the *MYC* promoter from untreated SEM cells (purple) or following 30 min treatment with 1.5 % 1,6-hexanediol (green), mean of three biological replicates. Only the enhancer region is shown. Differential tracks show the change in profile in hexanediol-treated samples: pink bars show increases; blue bars show decreases. Overlay of strand-specific nascent RNA-seq data under untreated or hexanediol treatment. Dark gray bars show RNA levels in untreated cells; green bars show levels in hexanediol-treated cells, representative of three biological replicates. **e** Capture-C from the *BCL2* promoter, as in (d). **f** Capture-C traces at genes that are transcriptionally downregulated (orange), upregulated (red) or unaffected (gray) by 30 min hexanediol treatment. Purple line shows the profile in untreated cells; green line is from hexanediol-treated cells; ribbon shows +/−1 SD for three replicates. Vertical gray bar indicates the capture point for each gene. Horizontal bars show 10 kb region around BRD4 ChIP-seq peaks. Shading highlights effect of IBET treatment on promoter interaction frequency within that window: pink bars indicate statistically-significant increases; blue bars indicate decreases; gray bars indicate no significant difference (Holm-Bonferroni adjusted p-value <0.05, paired Mann-Whitney test). Scale bar shows 100 kb.

Despite the strong downregulation of *MYC* and *BCL2* by hexanediol (Fig 4c), enhancer-promoter interactions at both genes were clearly retained (Fig 4d, e), although, as with IBET treatment, there were subtle rearrangements. In particular, unlike with IBET treatment, there was a slight reduction in enhancer-promoter interactions at *BCL2* (Fig 4e), but this is a minimal effect compared to enhancer-promoter reductions we have detected at other loci following DOT1Li ^30^ (Fig 3c). Thus, using three different drug treatments (IBET, JQ1, hexanediol) to reduce BRD4 and Mediator binding at the *MYC* and *BCL2* enhancers, we found very little evidence for a loss of interaction with their cognate promoters despite the reduction in transcription.

Analysis of other gene promoters by Capture-C revealed a similar lack of changes in interaction frequency following hexanediol treatment (Fig 4f), regardless of the transcriptional change at the gene (Supplementary Fig 4c). Surprisingly, as with IBET treatment, the most common difference appeared to be a slight increase in contact frequency (Fig 4f). Notably, statistical analysis identified a strong correlation in the effects observed at BRD4 peaks for the four treatments used (IBET for 90 min or 24h, JQ1 for 90 min and hexanediol for 30 min), and anticorrelation with DOT1L inhibition (Supplementary Fig 4d). Thus, three distinct drug treatments disrupting BRD4 localization produced a similarly weak effect on enhancer-promoter interactions at the genes studied. From this we conclude that reduction of BRD4 and MED1 binding at enhancers can have a strong impact on transcription, but this is not sufficient to significantly disrupt enhancer-promoter interactions. This contrasts strongly with our past work where loss of H3K79me2/3 at KEEs causes both decreased transcription as well as reduced enhancer-promoter interactions ^30^ (Fig 3c, d).

### Cohesin/CTCF binding patterns support a role in mediating a subset of enhancer-promoter interactions

Another mechanism that has been proposed to mediate interaction between promoters and enhancers is the loop extrusion model, which is also used to explain the generation of higher order chromatin structures. In this model, a loop of chromatin is fed through cohesin, until it is paused by two CTCF molecules bound in a convergent orientation ^9,10^. In SEM cells, CTCF and RAD21 show a strong positive correlation at ATAC peaks (Fig 1b), suggesting that all or most of these CTCF binding sites are competent to enrich or stabilize RAD21 association with chromatin. It is unclear whether loop extrusion may contribute to the increased local interactions between promoters and enhancers, although recent work has suggested this possibility ^21,22^. In support of this idea, CTCF and RAD21 can be observed at many enhancers and promoters, as well as non-enhancer/promoter ATAC peaks in SEM cells (Fig 1a). Further, the binding of CTCF or RAD21 at the *MYC* or *BCL2* promoter or enhancer is mostly unperturbed by IBET or hexanediol treatment (Supplementary Fig 5a, b), correlating with the maintenance of enhancer-promoter interactions.

To investigate whether cohesin/CTCF binding is a plausible mechanism to mediate enhancer-promoter contact, we compared the ChIP-seq profiles of these proteins to our Capture-C promoter interaction profiles. At *MYC* both the promoter and enhancer regions are associated with several closely-spaced CTCF/RAD21 peaks (Fig 5a). Strikingly, the promoter CTCF-bound motifs are oriented towards the enhancers, and the enhancer binding sites are oriented towards the promoter (Fig 5a, blue triangles), suggesting that any pairing of these would produce a convergent CTCF dimer, consistent with cohesin-mediated DNA looping ^10,55,56^. Indeed, the promoter CTCF sites have previously been shown to play a role in interaction with distinct enhancer regions in different cancer cell lines ^23^. At the *MYC* enhancer, the proximal domain is bounded by a pair of CTCF/RAD21 binding sites, and the major peak of the distal region is marked by two peaks (Fig 5a). Given that these all overlap with key points in the promoter interaction profile, this indicates that there may be multiple opportunities to stabilize contacts with the promoter via this mechanism.

**Figure 5.**
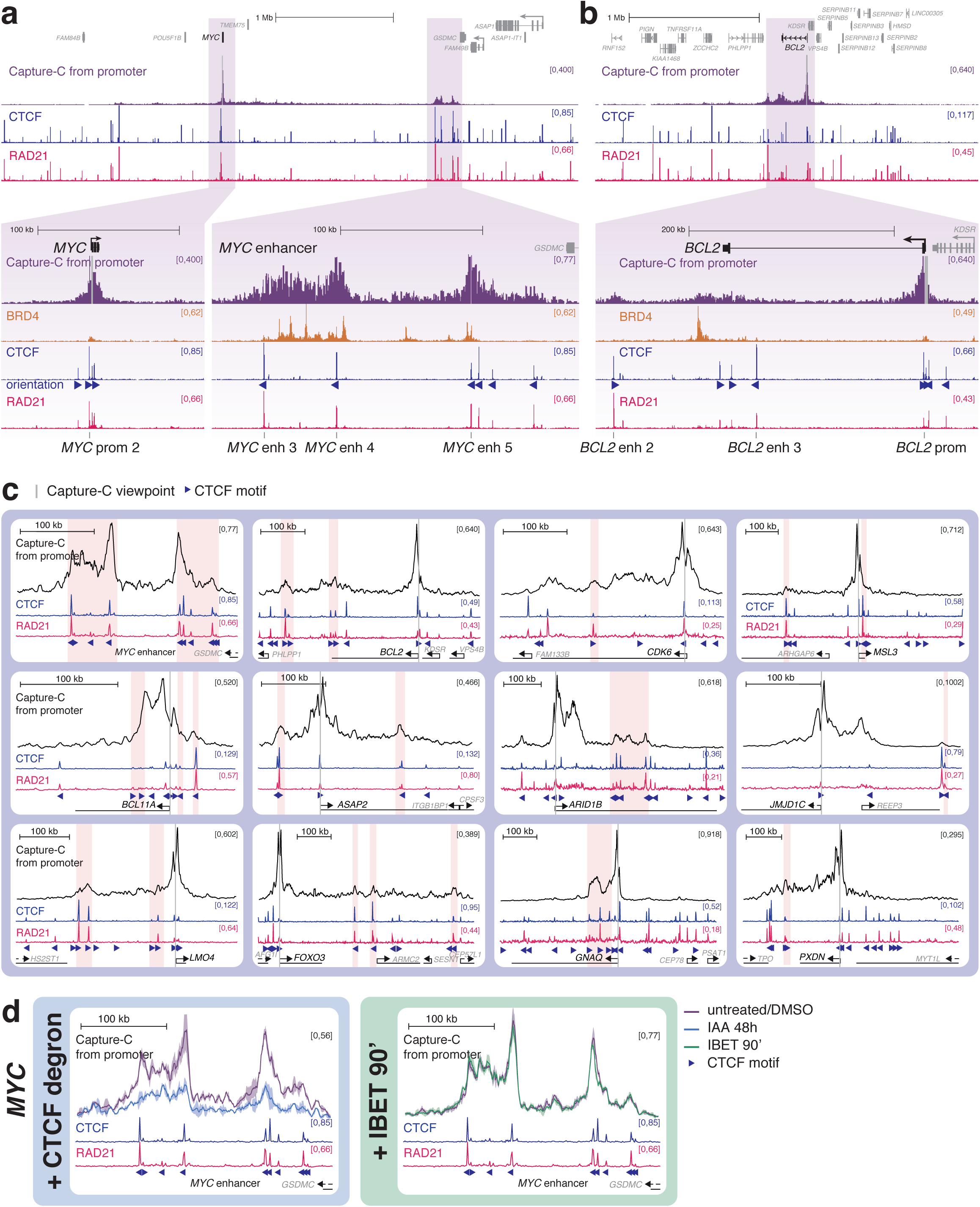
CTCF and RAD21 may be responsible for mediating enhancer-promoter interactions at *MYC* and *BCL2*. **a** Capture-C and ChIP-seq for BRD4, CTCF and RAD21 at the *MYC* gene and enhancer region. Capture-C was conducted using the *MYC* promoter as the viewpoint, indicated by a vertical gray bar, mean of three biological replicates. Orientation of CTCF motifs at peaks is indicated by triangles. Locations of primers used for CTCF/RAD21 ChIP-qPCR (see Supplementary Fig 5) are shown at the bottom of the figure. **b** Capture-C, ChIP-seq and ATAC-seq data at *BCL2*, as in (a). **c** Capture-C traces from untreated cells (black line; mean of three replicates) and ChIP-seq for CTCF (blue) and RAD21 (pink). Vertical gray bar indicates the capture point for each gene. Orientation of CTCF motifs at peaks is indicated by triangles. Scale bars show 100 kb. Shading shows CTCF/RAD21 peaks that overlap with enriched frequency of interaction (visually determined), with CTCF motifs oriented towards (pink) or away from (blue) the promoter. **d** *Left*: Capture-C profile from the *MYC* promoter showing the *MYC* enhancer in cells with AID-tagged CTCF, either untreated (purple line) or treated with indole-3-acetic acid (IAA) for 48h (blue line), which targets CTCF for degradation. Data are replotted from Hyle et al (2019), mean of two independent clones. *Right*: Capture-C profile from the *MYC* promoter showing the *MYC* enhancer in cells treated with DMSO (purple line) or IBET (green line) for 90 min, mean of three biological replicates. CTCF (blue) and RAD21 (pink) ChIP-seq tracks and CTCF motif orientations (triangles) are shown.

As at *MYC*, there are multiple CTCF/RAD21 peaks at the promoter of *BCL2*, and a clear convergent peak at the distal interaction region (which is not marked with BET proteins or other enhancer features) (Fig 5b; overlapping with the *BCL2* enh 2 primer pair; Fig 1d). There are also two CTCF sites (convergent with the promoter) which overlap with the broad interaction domain at the enhancer, although notably these CTCF sites occupy a distinct region to the peak of BRD4 and do not fully correlate with the interacting region. This suggests that additional or alternative factors to CTCF may facilitate contact between the *BCL2* enhancer and promoter.

We expanded this analysis to include other enhancer-associated genes for which we had promoter interaction data. Many RAD21/CTCF peaks within the analyzed domains were not associated with interactions with the target promoter (e.g. between the *MYC* promoter and enhancer, Fig 5a), but may be involved in mediating other DNA-DNA contacts. However, we identified many instances of promoter-oriented RAD21/CTCF peaks overlapping with enhancer-promoter interactions (Fig 5c, pink highlights).

Our data suggest that at a subset of enhancers, CTCF and cohesin may be partly responsible for facilitating enhancer-promoter interactions. Recent work in SEM cells, the cell line studied here, used Capture-C following CTCF degradation to test the effect of CTCF loss on the interaction between the *MYC* promoter and enhancer ^22^. Consistent with the model that CTCF/RAD21 binding stabilizes enhancer-promoter interactions, loss of CTCF was found to reduce both *MYC* expression and interactions between the *MYC* enhancer and promoter ^22^. Reanalysis of these data using the same approach as for our data contrasts the dramatic decreases in interaction observed with CTCF degradation with the minor changes observed following IBET inhibition (Fig 5d). These results argue that, at least at *MYC*, the loop extrusion model can explain enhancer-promoter contact and may be important for gene expression.

## Discussion

Maintenance of enhancer-promoter looping is thought to be crucial for gene activation, but emerging evidence and the data presented here suggest that enhancer-promoter contact and gene activation may be partially separable events. Importantly, physical interaction with the promoter may not be necessary for all enhancers ^5,6^, although it appears to be a requisite for most. At the same time, enhancer-promoter looping alone is not sufficient for activation, as enhancer-promoter contacts have been observed in the absence of transcription ^57,58^. Similarly, we show here that transcription can be disrupted with minimal changes in enhancer-promoter interaction frequency, as has been observed at the β-globin locus ^59^. In contrast, artificially stabilizing enhancer-promoter loops can activate transcription ^24,60–64^, indicating that the conversion of unproductive enhancer-promoter contacts to a functional complex may be dependent on the presence of additional factors.

These data suggest a model whereby stabilization of enhancer-promoter loops is a necessary but not sufficient precondition for gene activation, and the protein complexes that facilitate looping are not sufficient to directly promote gene expression. A distinct, functionally separable, stage of gene activation follows, where enhancer-associated factors interact with the promoter, producing transcriptional upregulation. Indeed, enhancer-promoter loop structures likely create an opportunity for contact between factors at these two loci. A number of enhancer-associated factors, including BRD4 and Mediator, have been observed to form phase-separated condensates *in vivo* ^2,34^, and we found that disruption of BRD4 and Mediator binding (under conditions that inhibit phase condensate formation) had appreciable effects on transcription. This suggests that there may be a role for the low-affinity interactions that can drive phase condensation in facilitating functional interactions between enhancers and promoters, particularly at regions of high activator density, such as super-enhancers, for example by stabilizing the binding of RNA polymerase at the promoter ^34,65,66^.

Recent models have proposed that the low-affinity interactions that drive phase condensation may be sufficient for both enhancer-promoter colocalization as well as promoting transcriptional activation ^3,67^. Computational simulation has suggested that formation of these structures may promote long-range chromatin interactions ^3^, and BRD4 is capable of driving clustering of acetylated chromatin *in vitro* ^68^. In support of this, BRD4 intrinsically disordered regions (IDRs) targeted to telomeric sequences appear to bring loci together ^35^ and dissolution of phase condensate structures prevents the estrogen-induced colocalization of enhancers^69^. However, it is unclear how closely these observations represent physiological enhancer-promoter interactions, or whether these results are representative of mechanisms functioning generally at most enhancers. Our data demonstrate that these low affinity interactions are not necessary for the maintenance of enhancer-promoter contacts, as reduction of BRD4 chromatin binding had no effect on promoter interaction profiles. A similar lack of effects was recently observed, albeit at lower resolution by Hi-C, in Mediator mutant mouse embryonic stem cells (mESCs) ^21^. Importantly, the high resolution and sensitivity of Capture-C confirms the lack of even subtle changes in enhancer-promoter contacts in our experiments, for example localized to BRD4 binding sites.

We note that, although our drug treatments reduced BRD4 and Mediator binding, it was not completely abolished, as detected by ChIP. This raises the possibility of a threshold effect in enhancer function. That is, high levels of BET/Mediator are needed for transcription, but low levels of BET/Mediator binding may be sufficient to maintain enhancer promoter interactions (Fig 6). Arguing against this model, the more common change we observed when BET/Mediator binding was reduced was actually a slight increase in enhancer-promoter interaction frequencies. This behavior is similar to that observed in a Mediator mutant cell line ^21^, suggesting that it may be a genuine consequence of Mediator loss. It is possible that these structural changes are an indirect effect of the transcriptional disruption following BRD4/Mediator loss, as has been observed before ^70,71^, although there was no correlation with transcriptional response.

**Figure 6.**
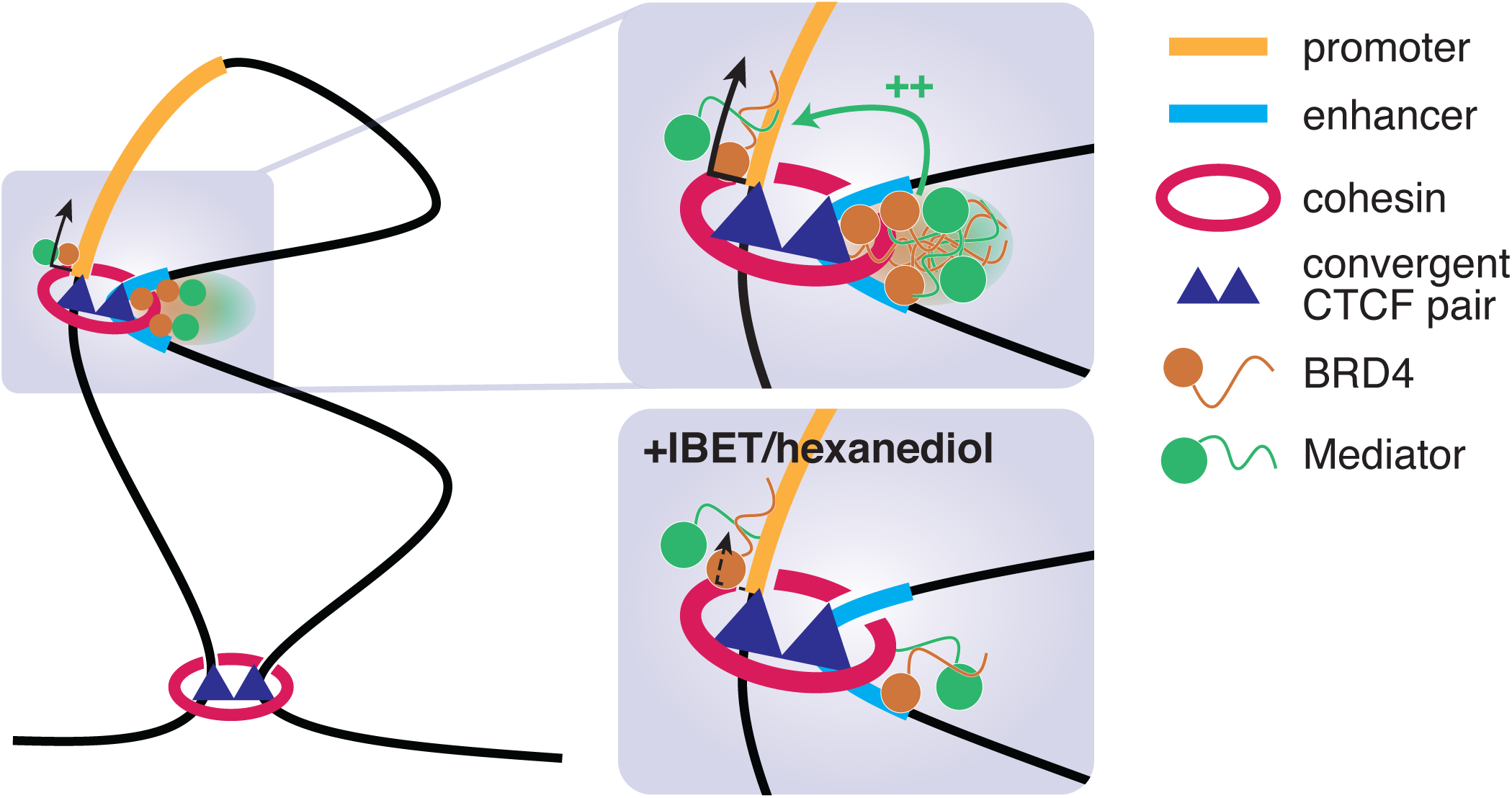
Model for enhancer-promoter interaction. Higher order chromatin boundaries are maintained by cohesin loops associated with convergent CTCF dimers. Within a domain, many enhancer-promoter contacts are associated with RAD21/CTCF peaks, and we suggest that similar cohesin loops are required for a subset of these interactions. At some enhancers (for example super-enhancers) a high concentration of factors such as BRD4 and mediator drive the formation of phase condensates, and these may increase interactions with factors at the promoter, held nearby by cohesin loops. These interactions may be required to activate or increase transcription from the promoter. Upon addition of IBET or 1,6-hexanediol, phase condensates are dissolved and BRD4 and mediator binding is reduced at the enhancer, disrupting interaction with factors at the promoter. This disrupts gene expression, but does not affect enhancer-promoter proximity as the two loci remain held together by cohesin.

What, then, is responsible for maintaining enhancer-promoter interactions? We favor a role for cohesin and CTCF, at least at a subset of genes, potentially in combination with low levels of BRD4 and Mediator (Fig 6) or other activators such as transcription factors. Whilst the loop extrusion model is widely accepted in the maintenance of higher order domain structure, a function at enhancers is less clear ^67^. Depletion of cohesin or its loader NIPBL disrupts interactions between promoters and distal enhancers ^21,72,73^, although this may be an indirect effect through loss of TAD boundaries rather than physical contact with enhancers. As has been reported ^23^, in our analysis we observed an enrichment for CTCF/RAD21 binding at promoter-interacting loci, consistent with a direct role for loop extrusion in mediating enhancer-promoter interactions. Indeed, loss of Rad21 results in decreased enhancer-promoter contacts and transcriptional downregulation at *Sik1* and *Elf3* in mESCs ^72^. Similarly, enhancer-promoter interactions at *MYC* are dependent on CTCF ^22,23^. However, it is possible that this gene is unusual, as long distance enhancer-promoter interactions over 1 Mb are not common. Strikingly, the majority of genes rapidly downregulated following CTCF degradation in mESCs show CTCF binding at the promoter, although this effect was proposed to be a result of a looping-independent function of CTCF in transcription ^74^.

One complicating aspect of the role for cohesin in enhancer-promoter contacts is the fact that disruption of loop extrusion, either by loss of cohesin itself, its loader NIPBL or CTCF, does not have widespread effects on gene expression ^22,73–76^. This suggests, assuming that the majority of enhancer-promoter interactions are productive, that cohesin is not essential for most enhancer function. However, given that our current understanding of enhancer function remains incomplete, this point alone is insufficient to rule out a role for loop extrusion in linking at least a subset of enhancers to promoters.

It is likely that multiple mechanisms exist to facilitate enhancer-promoter interactions at different genes, and may function at least partly redundantly. For example, deletion of the sole CTCF site in the *Sox2* super-enhancer in mESCs reduces, but does not abolish, contact with the promoter ^55^. Indeed, in our analysis, whilst many of the promoter interaction sites overlap with correctly oriented CTCF sites, there are also many sites of interaction that lack an obvious peak of CTCF binding (e.g. at *ARID1B*). We also observe broad regions of interaction that are bookended by peaks of CTCF/RAD21 (e.g. at the *MYC* enhancer), which suggests that, whilst CTCF and cohesin may define the borders of interaction, additional factors may play a role in more local contacts with the promoter. H3K4me1, a mark of enhancers, has itself been found to interact with cohesin ^29^. Transcription factors are another plausible anchor. Degradation of Oct4 (Pou5f1), but not Nanog, in mESCs results in a loss of Rad21 association with TF binding sites ^72^, arguing that specific TFs may be able to recruit or stabilize cohesin, potentially directing enhancer-promoter interactions. Additional structural proteins may also be important, for example YY1 ^24^. Mediator itself has been suggested to physically interact with cohesin ^77^.

There may also be a role for noncoding RNA in enhancer-promoter interactions ^25,26,78–80^. Notably, whilst mRNA has been shown to direct the formation of phase condensate compartments in the cytoplasm ^81^ and eRNAs have been proposed to play a similar role in the formation of enhancer-promoter complexes ^25,26^ our results following dissolution of phase condensates with hexanediol treatment argue that these interactions are not sufficient for maintaining contact at many genes. However, RNA may play other roles in directing enhancer-promoter interactions, for example by recruitment of cohesin ^79,80^.

Our results show that BRD4 and Mediator play a key role in the transcription of many genes, but they achieve this mainly via a functional rather than structural role. The high levels of these proteins at enhancers relative to promoters argues that contact between these loci is likely important for expression of many genes, but that this interaction functions primarily to enrich the local concentration of enhancer-bound factors at the promoter. Similarly, whilst the formation of phase condensates appears to be important for the transcription of many (but not all) genes, this is likely a mechanism to concentrate key transcription-related proteins at the enhancer-promoter complex, and a loss of these structures has little or no effect on enhancer-promoter looping. Physical contact between promoter and enhancer is not, *per se*, sufficient for transcription, and is not dependent on high levels of BRD4 or Mediator.

## Methods

### Cell culture and cell lines

SEM (an MLL-AF4 B-ALL cell line) ^82^ were purchased from DSMZ (www.cell-lines.de) and cultured in IMDM with 10% FBS and Glutamax. For drug treatments cells were diluted to 5 ×10^5^ cells/ml. IBET-151 was used at a final concentration of 1 µM, JQ1 at 1 µM and 1,6-hexanediol at 1.5 % (w/v).

### Chromatin Immunoprecipitation

ChIP-qPCR and ChIP-seq experiments were conducted as previously described ^49,83^. Briefly, double-fixed samples (2 mM disuccinimidyl glutarate (Sigma) for 30 min followed by 1 % formaldehyde (Sigma) for 30 min) were sonicated in batches of 10^7^ cells using a Covaris (Woburn, MA) following the manufacturer’s recommendations. Antibodies used for ChIP are detailed in Supplementary Table 3. Antibody-chromatin complexes were isolated using a 1:1 mixture of magnetic Protein A- and Protein G-dynabeads (ThermoFisher Scientific) and washed three times with a solution of 50 mM HEPES-KOH, pH 7.6, 500 mM LiCl, 1 mM EDTA, 1% NP-40, and 0.7% Na deoxycholate. Following a Tris-EDTA wash, samples were eluted with 50 mM Tris-HCl, pH 8.0, 10 mM EDTA and 1 % SDS, then treated with RNase A and proteinase K. DNA was purified using a PCR purification kit (Qiagen). For ChIP-qPCR, DNA was quantified relative to input chromatin, using primers listed in Supplementary Table 4. For ChIP-seq, DNA libraries were generated using the NEBNext Ultra II DNA Library Preparation kit (NEB). Samples were sequenced by 40 bp paired-end sequencing using a NextSeq 500 (Illumina).

### ChIP-seq bioinformatic analysis

Quality control of FASTQ reads, alignments, PCR duplicate filtering, blacklisted region filtering and UCSC data hub generation were performed using an in-house pipeline (https://github.com/Hughes-Genome-Group/NGseqBasic/releases). Briefly, QC was checked with fastQC (http://www.bioinformatics.babraham.ac.uk/projects/fastqc), then reads were mapped against the human genome assembly (hg19) using Bowtie ^84^. Unmapped reads were trimmed with trim_galore (https://www.bioinformatics.babraham.ac.uk/projects/trim_galore/) and remapped. Short unmapped reads from this step were combined using Flash and mapped again. PCR duplicates were removed with samtools rmdup ^85^, and any reads mapping to Duke blacklisted regions (UCSC) were removed with bedtools. Sequence tag (read) directories were generated from the sam files with the Homer tool makeTagDirectory ^86^. The command makeBigWig.pl was used to generate bigwig files for visualization in UCSC, normalizing tag counts to tags per 10^7^. Peaks were called using the Homer tool findpeaks.pl, with the input track provided for background correction, using -style histone or -style factor options to call peaks in histone modification or transcription factor datasets, respectively. Metagene profiles were generated using the Homer tool annotatePeaks.pl. Heatmaps were drawn using the R package heatmap3. CTCF motif orientations were assigned using the FIMO function of the MEME Suite ^87^.

### qRT-PCR

Total RNA was extracted and DNase I-treated from 10^6^ cell pellets using the RNeasy Mini kit (Qiagen). RNA was reverse-transcribed using SuperScript III (ThermoFisher Scientific) with random hexamer primers, then quantified using SyBr Green or TaqMan qPCR (see Supplementary Table 4 for primers). Gene expression was normalized to mature mRNA levels of the housekeeping gene *YWHAZ*.

### Nascent RNA-seq

Nascent RNA-seq extraction and purification was conducted as previously described ^49^. Briefly, 10^8^ SEM cells at 5×10^5^ cells/ml were treated with 500 µM 4-thiouridine (4-SU) for 1h (IBET treatments) or 30 min (hexanediol treatment), comprising the end of the drug treatment time (i.e. 30 min IBET treatment before 4-SU addition for 1 h, giving 90 min total IBET treatment time). Pelleted cells were lysed with Trizol (ThermoFisher Scientific) and total RNA was precipitated and DNase I-treated. 4-SU-incorporated RNA was biotinylated with EZ-link Biotin-HPDP (ThermoFisher Scientific) and purified with Streptavidin bead pull-down (Miltenyi). DNA libraries were generated from RNA using the NEBNext Ultra Directional RNA Library Preparation kit (NEB). Samples were sequenced by 75 bp paired-end sequencing using a NextSeq 500 (Illumina).

### RNA-seq bioinformatic analysis

Following QC analysis with fastQC (http://www.bioinformatics.babraham.ac.uk/projects/fastqc) reads were aligned against the human genome assembly (hg19) using STAR ^88^. Duplicate reads were removed using the picard command MarkDuplicates.jar (http://broadinstitute.github.io/picard). Gene expression levels were quantified as read counts using the featureCounts function from the Subread package with default parameters ^89^. The read counts were used to identify differential gene expression between conditions and generate RPKM values using the edgeR package ^90^. Genes were considered differentially expressed if they had an adjusted p-value (FDR) of less than 0.05. Strand-specific RNA-seq was visualized on UCSC using the bam file as input for homer commands makeTagDirectory (with options -flip and -sspe) and makeMultiWigHub.pl (with option -strand separate).

### Capture-C

Next-generation Capture-C was performed as previously described ^18^. Briefly, 2×110^7^ fixed SEM cell nuclei were digested with DpnII and used to generate a 3C library. Libraries were sonicated to a fragment size of 200 bp and Illumina paired-end sequencing adaptors (NEB) were added, using Herculase II (Agilent) to amplify the DNA. Indexing was performed in duplicate to maintain library complexity, with libraries pooled after indexing. Previously-designed Capture-C probes ^30^ targeting promoters or enhancers (Supplementary Table 1) were used to enrich for target sequences with two successive rounds of hybridization, streptavidin bead pulldown (ThermoFisher Scientific), bead washes (Nimblegen SeqCap EZ) and PCR amplification (NimbleGen SeqCap EZ accessory kit v2). Captured DNA was sequenced by 150 bp paired-end sequencing using a NextSeq 500 (Illumina). Data analysis was performed using an in-house pipeline (https://github.com/Hughes-Genome-Group/CCseqBasicF/releases). Capture-C promoter interactions overlapping with indicated ChIP-seq/ATAC-seq peaks were quantified for statistical analysis. Peaks outside of the bounds of Capture-C interaction domains (visually determined using UCSC genome browser) and those on *trans* chromosomes were removed from the analysis. Peaks within 10 kb of the Capture-C probe hybridization site were also removed. Holm-Bonferroni adjusted p-values for each peak were calculated by comparing all of the normalized read counts for each DpnII fragment and all replicates within a peak using a paired Mann-Whitney test for the two treatment conditions.

### Statistical analysis

Statistical analyses used and sample sizes are indicated in figure legends; n numbers refer to independent experiments. All tests were conducted two-tailed, all correlation analyses were conducted using the Pearson method.

### Data availability

All high throughput data has been deposited in the Gene Expression Omnibus (GEO) under the accession number GSExxxxxx. GEO accession numbers for datasets used from previous publications can be found in Supplementary Table 5.

### Materials and correspondence

Correspondence and material requests should be addressed to T.A.M.

## Acknowledgements

T.A.M., N.T.C., E.B., L.G., R.T., J.K., and M.T. were funded by Medical Research Council (MRC, UK) Molecular Haematology Unit grant MC_UU_12009/6 and MR/M003221/1. P.H. and J.O.J.D. are funded by an MRC Clinician Scientist Award to J.O.J.D. (MR/R008108). P.F. was supported by the Medical Research Council (MR/N010051/1). The authors would like to thank Jill Brown for her very helpful comments on the manuscript.

## Author contributions

N.T.C., E.B. and T.A.M. conceived the experimental design; N.T.C., E.B., L.G., T.A.M., J.K. and M.T. carried out experiments; P.F. provided reagents; N.T.C., R.T. and E.R. analyzed and curated the data; N.T.C., E.B. and T.A.M. interpreted the data; P.H. and J.O.J.D. provided expertise; N.T.C. and T.A.M. wrote the manuscript; all authors contributed to reviewing and editing the manuscript; T.A.M. provided supervision and funding.

## Supplementary Tables

**Supplementary Table 1.** Capture-C probes used in this study

**Supplementary Table 2.** Nascent RNA-seq data following treatment with 1 µM IBET-151 for 90 min or 24h or 1.5 % 1,6-hexanediol for 30 min

**Supplementary Table 3.**
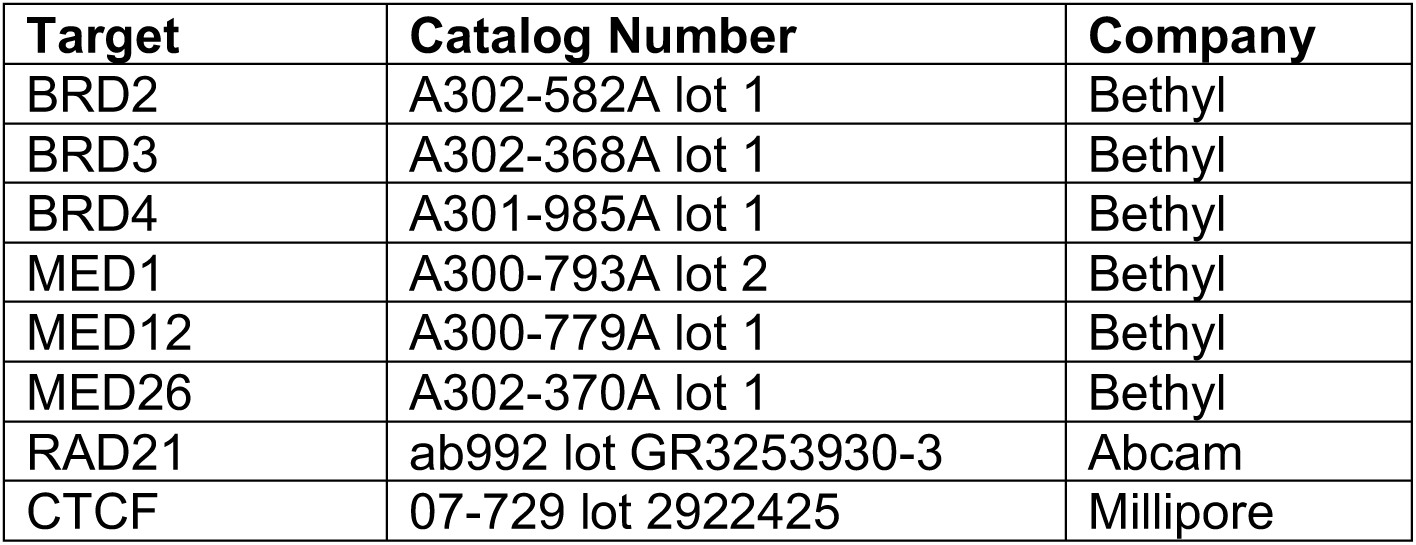
Antibodies used in this study

**Supplementary Table 4.**
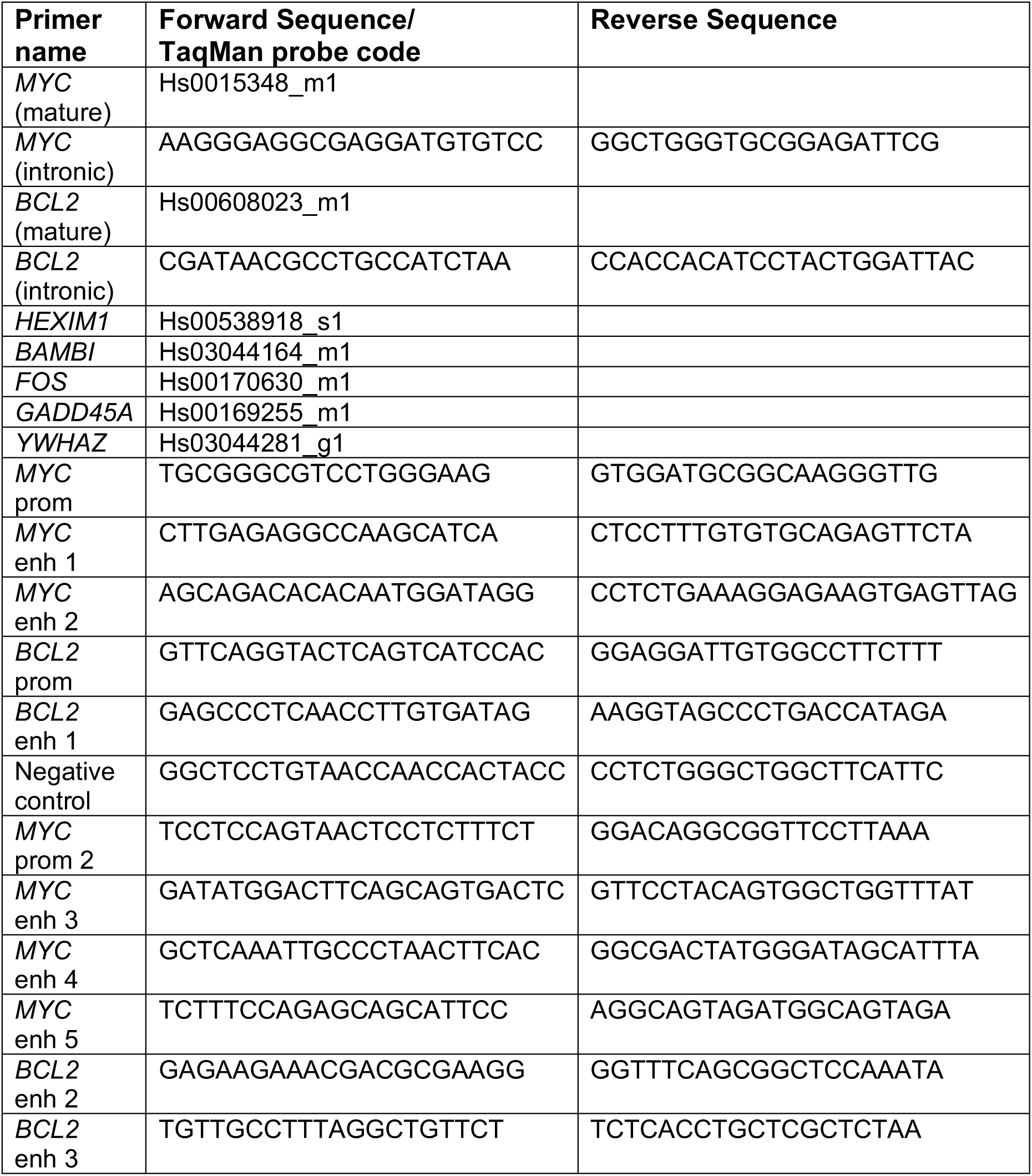
PCR Primers used in this study

**Supplementary Table 5.** Publicly available datasets used in this study

| **Data type** | **Cell type** | **Sample** | **GEO accession number** |
| --- | --- | --- | --- |
| ATAC-seq | SEM | Control | GSE117865 |
| Capture-C | SEM | DMSO 7d | GSE117865 |
| Capture-C | SEM | EPZ-5676 7d | GSE117865 |
| Capture-C | SEM CTCF-dTag Clone 27 | Untreated | GSE121257 |
| Capture-C | SEM CTCF-dTag Clone 27 | IAA 48h | GSE121257 |
| Capture-C | SEM CTCF-dTag Clone 35 | Untreated | GSE121257 |
| Capture-C | SEM CTCF-dTag Clone 35 | IAA 48h | GSE121257 |
| ChIP-seq | SEM | H3K4me1 | GSE74812 |
| ChIP-seq | SEM | H3K4me3 | GSE74812 |
| ChIP-seq | SEM | H3K27ac | GSE74812 |
| ChIP-seq | SEM | BRD4 | GSE83671 |
| ChIP-seq | SEM | MED1 | GSE83671 |
| ChIP-seq | SEM | CTCF | GSE117865 |
| ChIP-seq | SEM | ELF1 | GSE117865 |
| ChIP-seq | SEM | ERG | GSE117865 |
| ChIP-seq | SEM | FLI1 | GSE117865 |
| ChIP-seq | SEM | MYB | GSE117865 |
| ChIP-seq | SEM | RUNX1 | GSE42075 |
| ChIP-seq | SEM | RUNX2 | GSE117865 |
| ChIP-seq | SEM | SPI1 | GSE117865 |

## Supplementary Figure Legends

**Supplementary Figure 1.**
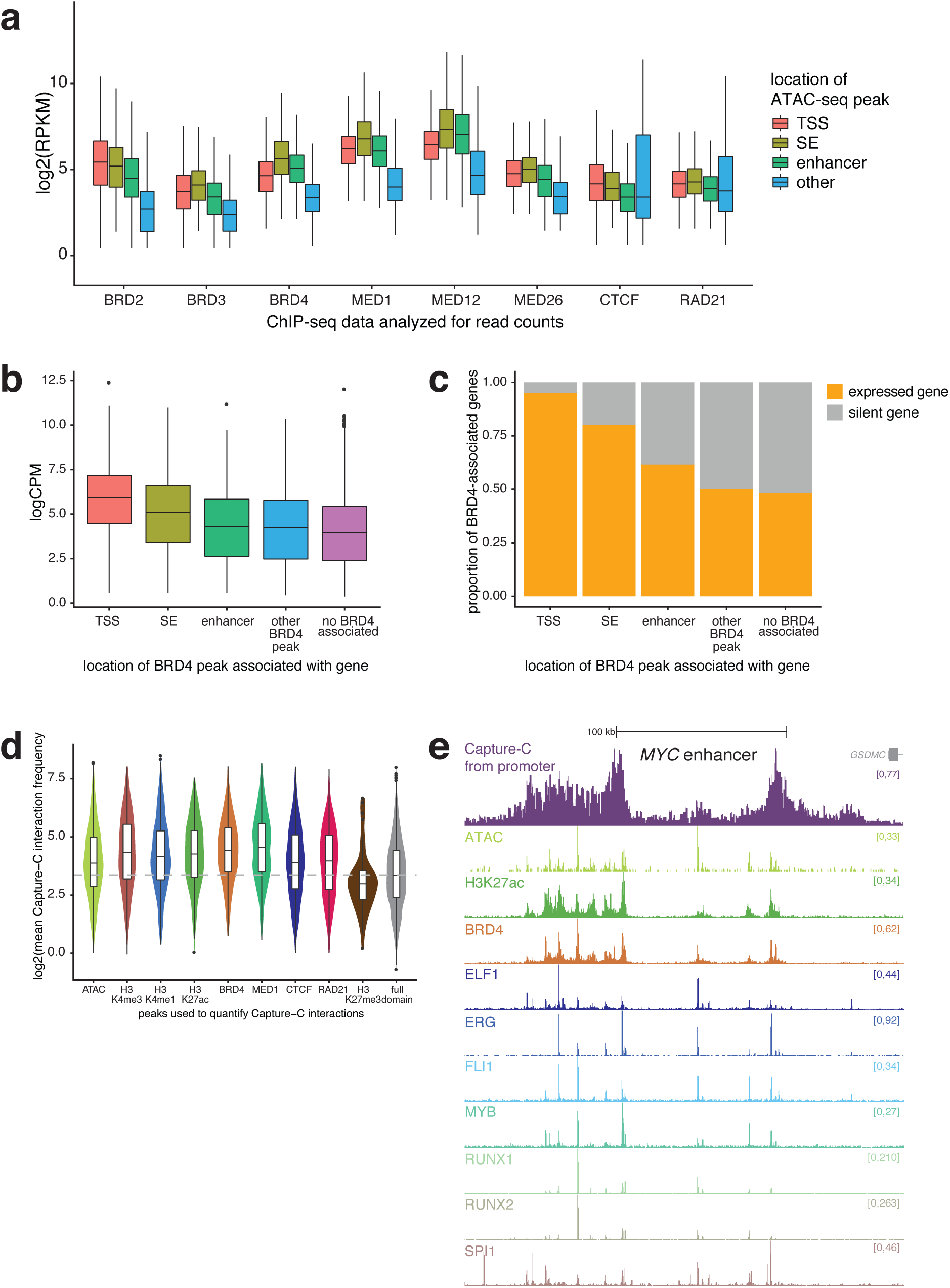
**a** Boxplot showing levels of BET proteins, Mediator subunits, CTCF and RAD21 at ATAC peaks found at TSSs (red), within super-enhancers (SE; olive), within other enhancers (green) or at other sites (blue) in SEM cells. **b** Level of expression of genes (logCPM from nascent RNA-seq), classified based on the location of the nearest BRD4 peak. **c** Proportion of expressed genes, classified based on the location of the nearest BRD4 peak. **d** Frequency of promoter interactions at 10 kb regions flanking ATAC-seq/ChIP-seq peaks for the indicated antibodies. Boxplots show the median and IQR. Full domain: data for the entire analyzed regions divided into 10 kb bins. **e** Transcription factor ChIP-seq tracks at the *MYC* enhancer. Capture-C from the *MYC* promoter (mean of three replicates), ATAC-seq and H3K27ac and BRD4 ChIP-seq tracks are reproduced from Fig 1c for comparison.

**Supplementary Figure 2.**
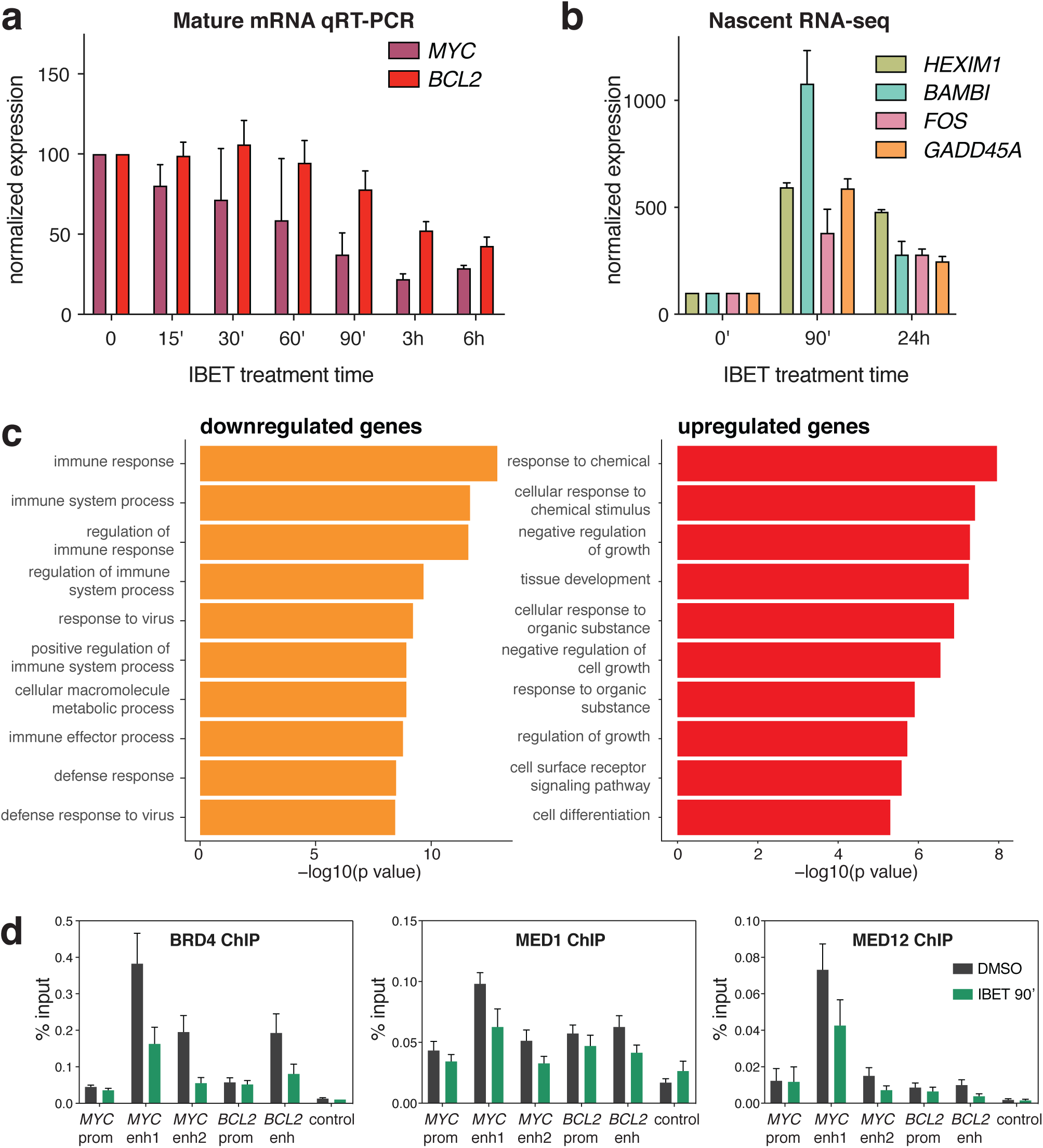
**a** qRT-PCR analysis of RNA levels following 1µM IBET treatment for the indicated times, using mature mRNA PCR primers. Values are normalized to *YWHAZ* mature mRNA levels, relative to DMSO treatment. Mean of three biological replicates, normalized to expression in DMSO; error bars show SEM. **b** Quantification of nascent RNA-seq levels of upregulated genes shown in Fig 2a, following 90 min or 24h IBET treatment. Data are cpm-normalized, relative to expression levels under DMSO treatment. Mean of three biological replicates; error bars show SEM. **c** Top ten biological pathway GO terms most significantly enriched in down- and upregulated genes (logFC <−1 or >1, FDR <0.05) following 90 min IBET treatment. **d** ChIP-qPCR for BRD4, MED1 and MED12 following 90 min treatment with DMSO (black) or IBET (green). Mean of four biological replicates; error bars show SEM. Primer locations are shown in Fig 1c-d.

**Supplementary Figure 3.**
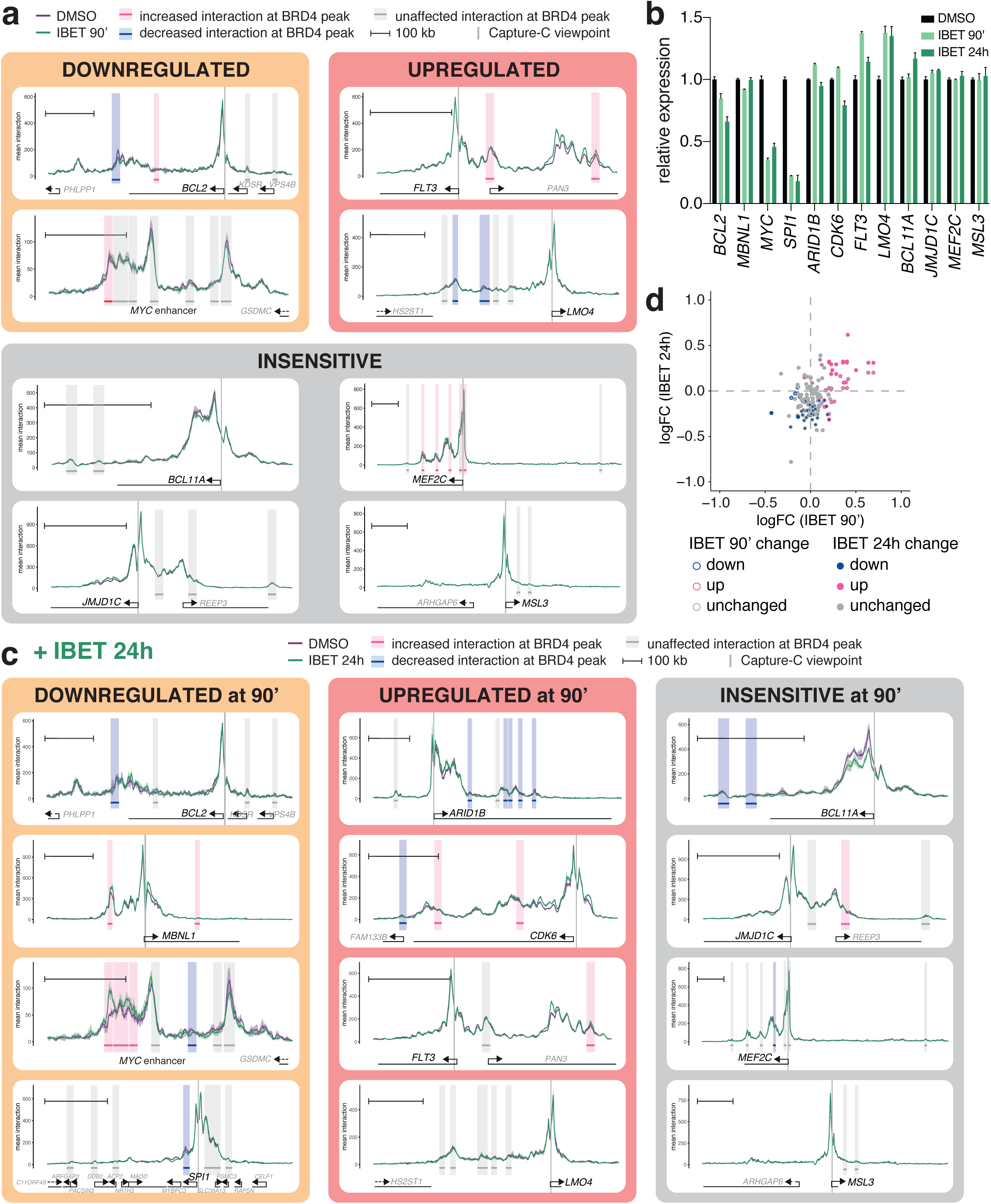
**a** Capture-C traces at genes which are transcriptionally downregulated, upregulated or unaffected following 90 min 1µM IBET treatment. Purple line shows the profile in DMSO-treated cells; green line is from cells treated with IBET for 90 min; ribbon shows +/−1 SD for three replicates. Vertical gray bar indicates the capture point for each gene. Horizontal bars show 10 kb region around BRD4 ChIP-seq peaks. Shading highlights effect of IBET 90 min treatment on promoter interaction frequency within that window: pink bars indicate statistically-significant increases; blue bars indicate decreases; gray bars indicate no significant difference (Holm-Bonferroni adjusted p-value <0.05, paired Mann-Whitney test). Scale bar shows 100 kb. **b** Quantification of nascent RNA-seq expression of genes shown in Fig 3c and Supplementary Fig 3a, following 90 min or 24h 1 µM IBET treatment. Data are cpm-normalized, relative to expression levels under DMSO treatment, mean of three biological replicates. **c** Capture-C traces at genes which are transcriptionally downregulated, upregulated or unaffected following 90 min IBET treatment. Purple line shows the profile in DMSO-treated cells; green line is from cells treated with IBET for 24h; ribbon shows +/−1 SD for three replicates. Vertical gray bar indicates the capture point for each gene. Horizontal bars show 10 kb region around BRD4 ChIP-seq peaks. Shading highlights effect of IBET 24h treatment on promoter interaction frequency within that window: pink bars indicate statistically-significant increases; blue bars indicate decreases; gray bars indicate no significant difference (Holm-Bonferroni adjusted p-value <0.05, paired Mann-Whitney test). Scale bar shows 100 kb. **d** Comparison of changes in Capture-C promoter interactions following 1µM IBET treatment for 90 min or 24h. Mean of three biological replicates. Outer color indicates the effect of 90 min IBET treatment on interaction at each BRD4 peak, inner color indicates the effect of 24h treatment. Blue: decreased interaction; pink: increased interaction; gray: no change in interaction (Holm-Bonferroni adjusted p-value <0.05, paired Mann-Whitney test).

**Supplementary Figure 4.**
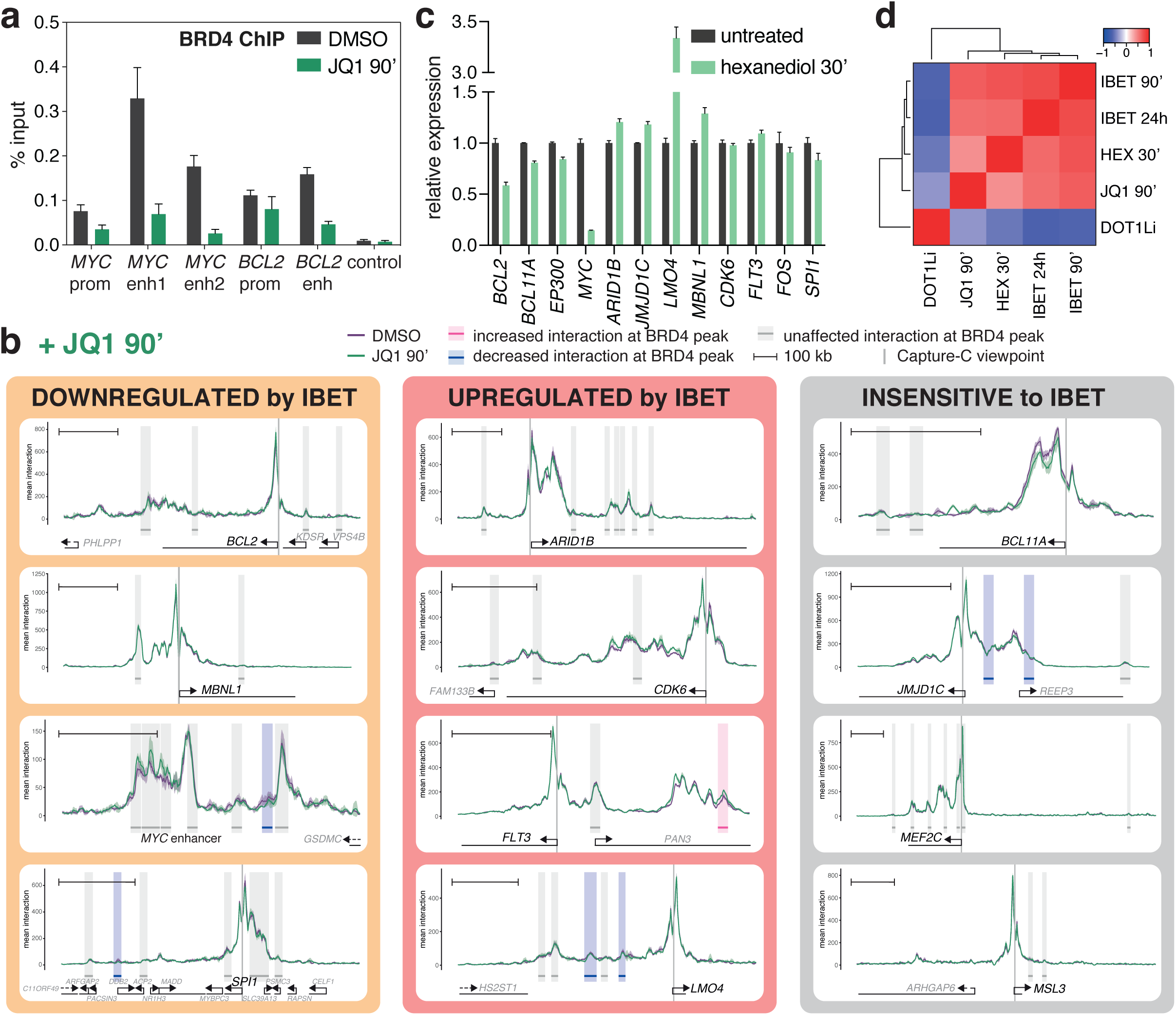
**a** ChIP-qPCR for BRD4 following 90 min treatment with DMSO (black) or 1 µM JQ1 (green). Mean of five biological replicates; error bars show SEM. Primer locations are shown in Fig 1c-d. **b** Capture-C traces at genes which are transcriptionally downregulated, upregulated or unaffected following 90 min IBET treatment. Purple line shows the profile in DMSO-treated cells; green line is from cells treated with JQ1 for 90 min; ribbon shows +/−1 SD for three replicates. Vertical gray bar indicates the capture point for each gene (in black, below). Horizontal bars show 10 kb region around BRD4 ChIP-seq peaks. Shading highlights effect of JQ1 treatment on promoter interaction frequency within that window: pink bars indicate statistically-significant increases; blue bars indicate decreases; gray bars indicate no significant difference (Holm-Bonferroni adjusted p-value <0.05, paired Mann-Whitney test). Scale bar shows 100 kb. **c** Quantification of nascent RNA-seq expression of genes shown in Fig 4f, following 30 min 1,6-hexanediol treatment. Data are cpm-normalized, relative to expression levels in untreated cells, mean of three biological replicates. **d** Pearson correlation of the changes in interaction frequency (logFC) between BRD4 peaks and promoters (10 kb windows) following the indicated treatments. Dendrogram shows hierarchical clustering of datasets.

**Supplementary Figure 5.**
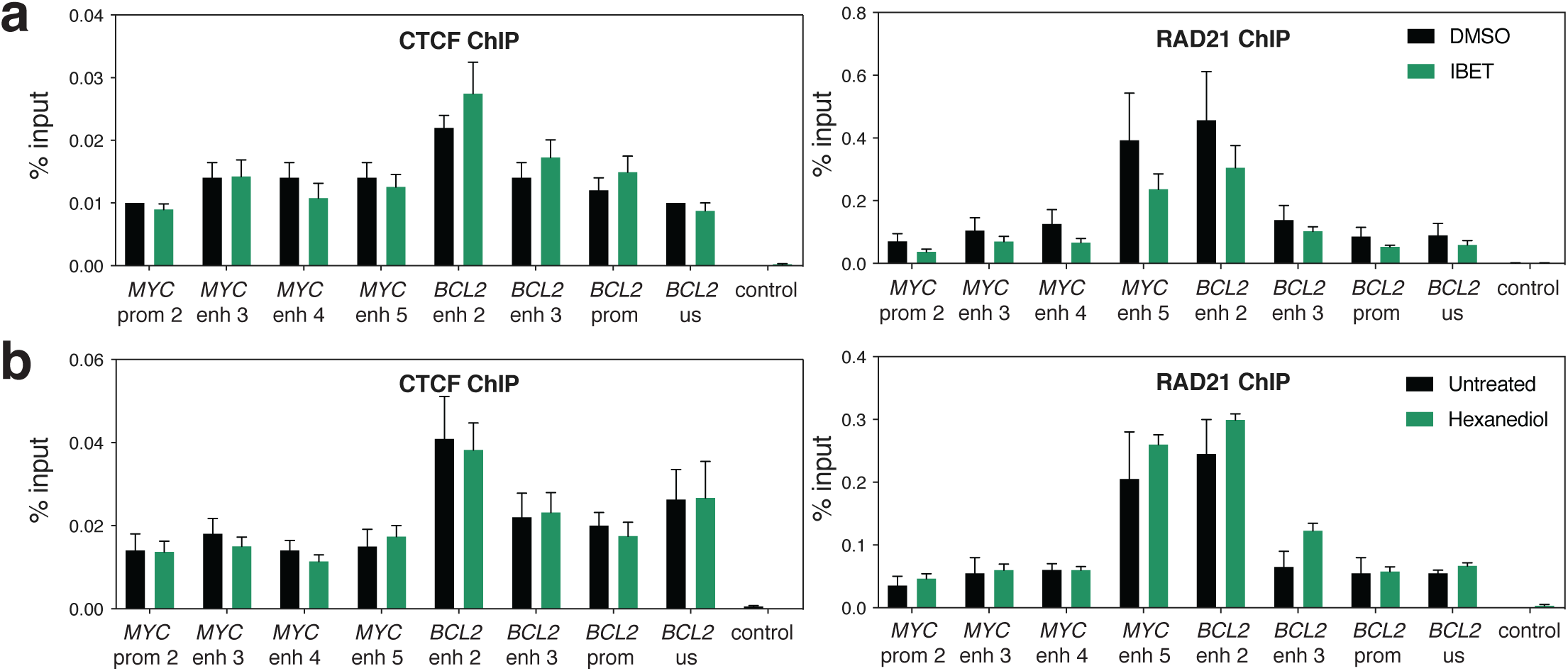
**a** ChIP-qPCR for CTCF and RAD21 in SEM cells following 90 min treatment with DMSO (black) or 1 µM IBET (green). Mean of four biological replicates; error bars show SEM. **b** ChIP-qPCR for CTCF and RAD21 from untreated SEM cells (black) or following 30 min treatment with 1.5 % 1,6-hexanediol (green). Mean of four biological replicates; error bars show SEM.

